# Immuno-phenotypes of Pancreatic Ductal Adenocarcinoma: Metaanalysis of transcriptional subtypes

**DOI:** 10.1101/198903

**Authors:** Ines de Santiago, Christopher Yau, Mark Middleton, Michael Dustin, Florian Markowetz, Shivan Sivakumar

## Abstract

Pancreatic ductal adenocarcinoma (PDAC) is the most common malignancy of the pancreas and has one of the highest mortality rates of any cancer type with a 5-year survival rate of < 5% and median overall survival of typically six months from diagnosis. Recent transcriptional studies of PDAC have provided several competing stratifications of the disease. However, the development of therapeutic strategies will depend on a unique and coherent classification of PDAC. Here, we use an integrative meta-analysis of four different PDAC gene expression studies to derive the consensus PDAC classification. Despite the fact that immunotherapies have yet to have an impact in treatment of PDAC, the gene expression signatures that stratify PDAC across studies are immunologic. We define these as “adaptive”, “innate” and “immune-exclusion” immunologic signatures, which are prognostic across independent cohorts. An appreciation of the immune composition of PDAC with prognostic significance is an opportunity to understand distinct immune escape mechanisms in development of the disease and design novel immune-oncology therapeutic strategies to overcome current barriers.

## Introduction

Despite on-going improved understanding of the genetics and molecular biology of Pancreatic Ductal Adenocarcinoma (PDAC), prognosis remains strikingly poor with five-year survival rate < 5% [1]. As surgical removal of the tumour remains the only curative treatment, the key to better outcomes currently depends entirely on early detection. Improving outcomes may be achieved through accurate subtyping upon detection to better tailor therapeutic strategies.

Using transcriptomics is an attractive option to sub-classify cancers. Gene expression profiles of tumours are a combined readout of the tumours’s genetic and epigenetic status, as well as the composition of other cells in the microenvironment that are sampled in the biopsy. The microenvironment includes both tumour stroma, host connective tissue cells that change in response to signals from the tumour cells, and innate and adaptive immune system cells [2, 3]. Several recent studies have described the transcriptomic landscape of PDAC and identified different subtypes with different clinical outcomes and drug sensitivities [4-7]. Table 1 summarises all the studies. Collisson et al proposed the first classification in 2011 by using unsupervised clustering methods on transcriptomic data [4]. This had identified three subtypes: classical, quasimesenchymal and exocrine-like, which have different prognoses and different responses to treatment. This classification has subsequently been challenged and revised, with newer classifications being put forward in 2015, 2016 and 2017 [5-7]. Moffitt *et al* use a deconvolution method to separate expression data based on stromal and normal gene profiles, which identified 4 groups: 2 stroma and 2 non-stroma tumour groups [5]. Moffitt subtyped PDAC ‘classical’ and ‘basal-like’ based on the pancreatic tumour cells and ‘normal stroma’ and ‘activated stroma’ based on the micro-dissected stoma cell population. The basal-like sub-type had the worst prognosis. Bailey *at al* took an integrative approach, which demonstrated 4 subtypes with different characterisations and different prognoses [6]. Bailey used non-negative matrix factorization and gene expression profiles to identify distinct subtypes of PDAC ‘squamous’, ‘pancreatic progenitor’, ‘immunogenic’ and ‘aberrantly differentiated endocrine exocrine’ (ADEX) [6]. Sivakumar *et al* used a master regulator approach based on the transcriptional effects of oncogenic *KRAS* to determine 3 subtypes that showed different prognoses, characterisations and treatment strategies[7]. Sivakumar et al., 2017, used master regulator analysis and community search and identified ‘Notch’ signalling, ‘Hedgehog/Wnt’ signalling, and ‘cell cycle’ as core processes of three distinct PDAC subtypes. A strong positive correlation between Hedgehog and Notch processes was observed meaning that activated Notch signalling was also present in the Hedgehog/Wnt group [7].

**Table 1:** Pancreatic ductal carcinoma and cancer subtyping signatures and studies used in this study

|  | Technique of discovery | Sub-type | Prognosis | Composition | Therapeutics |
| --- | --- | --- | --- | --- | --- |
| Collisson | Non-negative matrix factorization (NMF) analysis with consensus clustering | Classical | Good | N/A | Erlotinib |
|  |  | Exocrine | Average | N/A | Non-proposed |
|  |  | Quasi-mesenchymal | Poor | N/A | Gemcitabine |
| Moffitt | Non-negative matrix factorization (NMF) and digital separation | Tumour: classical | Average (median survival 19 months) | N/A | N/A |
|  |  | Tumour: basal-like | Poor (median survival 11 months) | N/A | Better response to adjuvant therapy |
|  |  | Stroma: normal | Good (median survival 20 months) | Stellate cells | N/A |
|  |  | Stroma: activated | Poor (median survival 15 months) | Macrophages | N/A |
| Bailey | Unsupervised clustering | Squamous | Poor | Macrophages | N/A |
|  |  | Pancreatic Progenitor | N/A | N/A | N/A |
|  |  | Immunogenic | N/A | B cells and T cells | Checkpoint blockade |
|  |  | ADEX | N/A |  |  |
| Sivakumar | Master regulator Analysis | Notch | Good | T cell infiltration | Immunotherapy |
|  |  | Cell cycle | Average | N/A | Adjuvant therapy |
|  |  | Hedgehog | Poor | Macrophages | N/A |

These studies suggest that it is possible to categorise PDAC through transcriptomics. However, this has not become the accepted clinical approach because of problems inherent with the approach – too many non-overlapping signatures, inadequate clinical relevance and poor mechanistic underpinning. It would thus be valuable to consolidate different stratification schemes into consensus subtypes of pancreatic cancer and develop reliable and robust biomarkers to better predict outcomes and select therapeutic strategies. In response to this unmet need, we have developed a consensus classification scheme from the four major transcriptomic PDAC sub-typing studies. PDAC has previously been shown previously to have different immune populations in its microenvironment as well as stroma [8]. In light of the tumour having a complex immune composition, the tumour microenvironment can be immunosuppressive or tolerant and this can be due to immune dysfunction [9, 10]. However, immunotherapy approaches have not yet had an impact on pancreatic cancer survival [11, 12]. Our meta-analysis reveals that the greatest prognostic value in independent cohorts could be better achieved through stratification by gene expression signatures associated with tumour infiltrating immune cells across different PDAC subtypes. Recognising the existence of different tumour escape mechanisms (and indeed phenotypes) in PDAC may guide distinct immunotherapeutic treatment plans and improve patient stratification for maximization of therapeutic effect.

## Results

### Consensus PDAC subtypes

We used published exemplar gene signature for data from Collison et al. 2011 and Moffitt et al. 2015 to cluster 242 PDAC primary tumour cases from the PACA-AU cohort (**Figure S1**). We further obtained and applied clustering labels identified by Bailey et al 2016 and Sivkumar et al 2017 also from the PACA-AU cohort. **Figure 1A** summarizes the workflow of our analysis.

**Figure 1.**
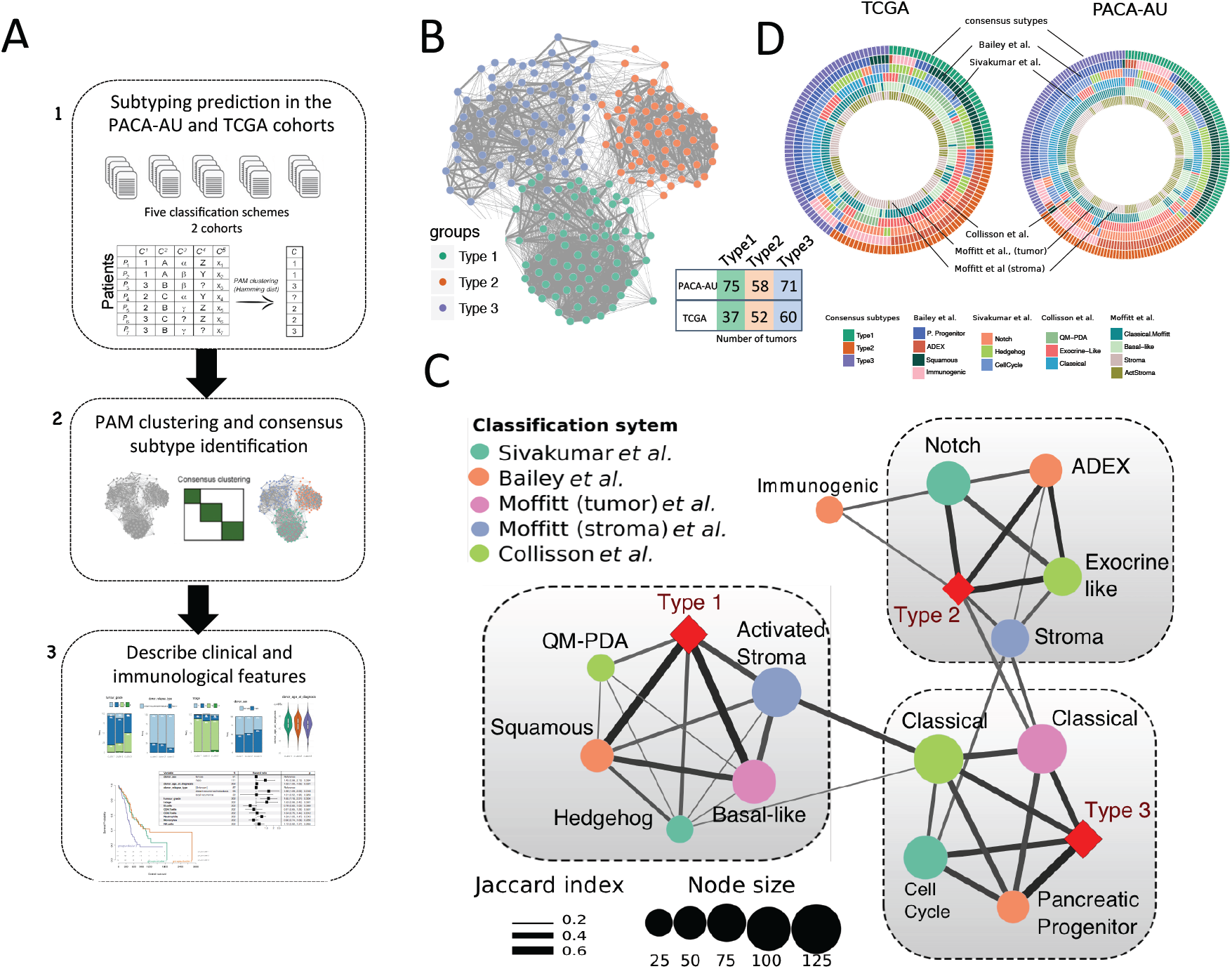
Identification of the consensus subtypes of PDAC. **(A)** Analytical workflow of the PDAC subtyping: (1) subtype classification using methodology from five different classification schemes; (2) concordance analysis of the five subtyping labels and application of PAM clustering algorithm to identify consensus clusters; (3) analysis of clinicopathological and immunophenotypes of PDAC consensus subtypes and identification of immune cell signatures with prognostic value in independent PDAC cohorts. **(B)** Patient similarity network. Each node represents a single patient sample in the PACA-AU cohort (n= 204). Network edges correspond to highly concordant (at least 5 of 6) subtyping calls between samples. Nodes are colored according to the three clusters identified from the PAM consensus clustering algorithm. **(C)** Circular heatmap representing sample overlap for consensus PDAC subtypes and mRNA subtypes from Bailey et al., Sivakumar et al., Collisson et al., or Moffitt et al. (from inside to outside, respectively). Significance of sample overlap was assessed with the hypergeometric test, adjusted p-values for each pairwise comparison are depicted in Supplementary Figure S4. **(D)** Association of consensus PDAC subtypes identified by PAM clustering (red refers to this study: Type 1, Type 2, Type 3) with tumour labels across five classification systems. Each node corresponds to a single subtype (circles are colored according to classification study; red diamonds correspond to the consensus subtypes). Edge width corresponds to the overlap between labels assessed by the Jaccard coefficient in the PACA-AU cohort, only significant edges are depicted (hypergeometric P ≤ 0.05. The three grey rectangles delineate clusters of tumour labels that overlap the three PDAC consensus groups (Type 1-3) with FDR adjusted P ≤ 0.05.

To identify a meaningful consensus of the multiple clustering labels we used consensus clustering and the partition around medoids (PAM) algorithm to cluster the 204 PDAC tumours according to the similarity of their labels and identified three robust consensus subtypes (**Figure 1B; Supplementary Figure S2**). Visualization of the similarity network between tumour labels revealed that consensus samples remained grouped between the three large primary network hubs (**Figure 1B**). We used 150 TCGA samples that were previously classified to perform the same consensus classification using the PAM algorithm in an independent cohort (Figure S3). Notably, the same three robust consensus subtypes were identified in the TCGA cohort (Figure 1C) with the three main consensus subtypes showing a stable and consistent pattern of co-associated labels (**Figure 1D; Supplementary Figure S4**). The Type 1 subtype cluster overlapped between the ‘basal-like’, ‘squamous’, ‘QM-PDA’ and ‘activated stroma’ subtypes (**Figure 1D**). The Type 2 subtype cluster, consisted of a statistically significant overlap between the ‘ADEX’, ‘exocrine-like’, ‘Notch’ and ‘normal-stroma’ subtypes. Finally, the subtype 3 cluster overlapped with the ‘classical’ subtypes identified by Moffitt and Collison studies and ‘cell-cycle’ group identified by Sivakumar et al., 2017 (overlaps were considered significant at hypergeometric FDR adjusted P ≤ 0.05 in both cohorts; **Figure 1D**).

### Clinicopathological characteristics of PDAC subtypes

Clinical features, including age, gender, tumour grade, TNM system tumour stage and tumour relapse, were statistically compared between the three clusters. Two-way contingency table analysis showed significant association between grade and cluster subtype, with subtype 1 tumours more likely to be grade 3 (Chi-square test P=0.002 PACA-AU cohort; **Figure 2A**) and subtype 2 tumours more likely to be grade 1 (Chi-square test P*=*0.033 for the TCGA cohort; **Figure 2B**). Tumour stage, relapse and gender did not correlate with the subtypes in any of the cohorts. Similarly, the average age of diagnosis was not significantly associated to any cluster subtype (**Figure 2A,B**; **Supplementary Table 1**).

**Figure 2:**
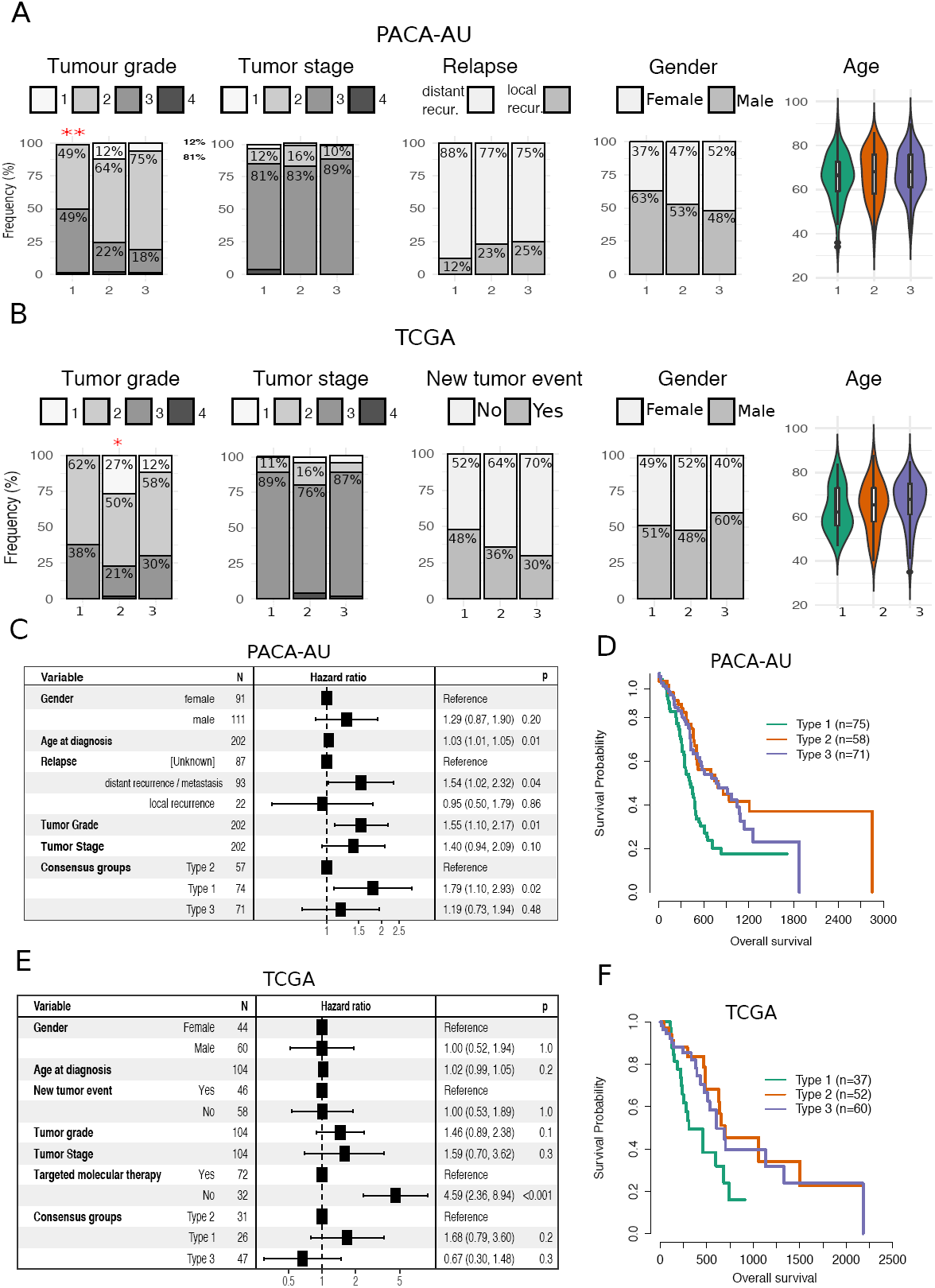
Clinicopathological and prognostic associations of consensus PDAC subtype groups. **(A,B)** Clinicopathological and prognostic associations of consensus PDAC subtype groups in the PACA-AU (A) and TCGA (B) cohorts. Distribution of histopathological grade; TNM system tumour stage at diagnosis, relapse type (PACA-AU) or new tumour event indicator (TCGA); gender; and age at diagnosis; across the three consensus subtypes, represented by the coloured violin plots subtype 1 (green); subtype 2, (orange); subtype 3 (violet). Red asterisks represent significant codes for Chi-square test p-values: *P <* 0.01 '**'; *P <* 0.05 '*'. (**C, E**) Multivariate Cox proportional hazards regression analysis for the PACA-AU (C) and TCGA (E) cohorts, with covariates including patient age at diagnosis, relapse, tumour stage (TMN system) and tumour grade. Squares represent the hazard ratio (HR) and the horizontal bars extend from the lower limit to the upper limit of the 95% confidence interval of the estimate of the hazard ratio. The plot also shows the number of considered events (N) and p-values (p) for the interaction between survival and any covariate. Detailed statistics are in **Supplementary Tables 1 and 2.** CI, confidence interval; HR, hazards ratio; p, Wald test p-value. (**D, F**) Prognostic value of subtype 1, 2 and 3 PDAC groups in the PACA-AU (D) and the TCGA (F) cohorts with Kaplan-Meier overall survival analysis.

To determine whether the PDAC clusters differed in outcome, we performed a Cox proportional hazards analysis. We observed differences in prognosis of different PDAC subtypes, with subtype 1 tumours associated to worse overall survival and higher HR (**Figure 2C-F; Supplementary Table 2**) in multivariate analyses, after adjustment for several clinicopathological features, including age, gender, tumour stage and grade. This difference was only statistically significant for PACA-AU cohort (subtype 1 vs subtype 2: P <0.05; HR=1.8; CI=[1.1-2.9]) and not in the TCGA cohort (subtype 1 vs subtype 2: P=0.2; HR = 1.7; CI=[0.8-3.6]). However we noticed a strong influence (*P* << 0.001; HR=4.6; CI=[2.4-8.9]) of ‘targeted therapy’ variable in overall survival (**Figure 2F**) and this might explain our inability to completely reproduce survival results observed in the PACA-AU cohort. Differential prognosis associated to the stroma type were also evident, as “classical” tumours with “normal” stroma subtypes had the best prognosis, while “basal-like” subtype tumours with “activated” stroma subtypes had the worse prognosis. Patients in the ‘Notch’ group of Sivakumar classification (enriched in subtype 2) also displayed the best overall survival rates [7].

### Immuno-phenotypes of Pancreatic Adenocarcinoma

Analysis of the tumour microenvironment has revealed that populations of tumour infiltrating immune cells have significant prognostic value in a variety of solid tumours [2, 13, 14]. Analysis in Melanoma, Breast Cancer and Colorectal Cancer has shown that tumour progression is characterized by distinct immune patterns [15-18] and that assessment of this ‘immunophenotype’ may provide better prognostic value beyond that predicted by traditional staging [19].

To further explore the composition of different immune infiltrates in the different tumour microenvironments of PDAC cancers we used single sample gene set enrichment analysis (ssGSEA) to score each tumour based on gene signatures representative of different cell types [20]. To test the consistency of the results we performed the analysis in the PACA-AU and the TCGA cohorts. We used four published signatures from independent studies [21-24] and focused on 7 immune cell types (NK cells, Neutrophils, Macrophages, Dendritic cells, CD4+ T-cells, CD8+ T-cells and B-cells). We observed that immune signatures are differentially enriched in the tumour microenvironment of PDAC subtypes (**Figure 3A**).

**Figure 3.**
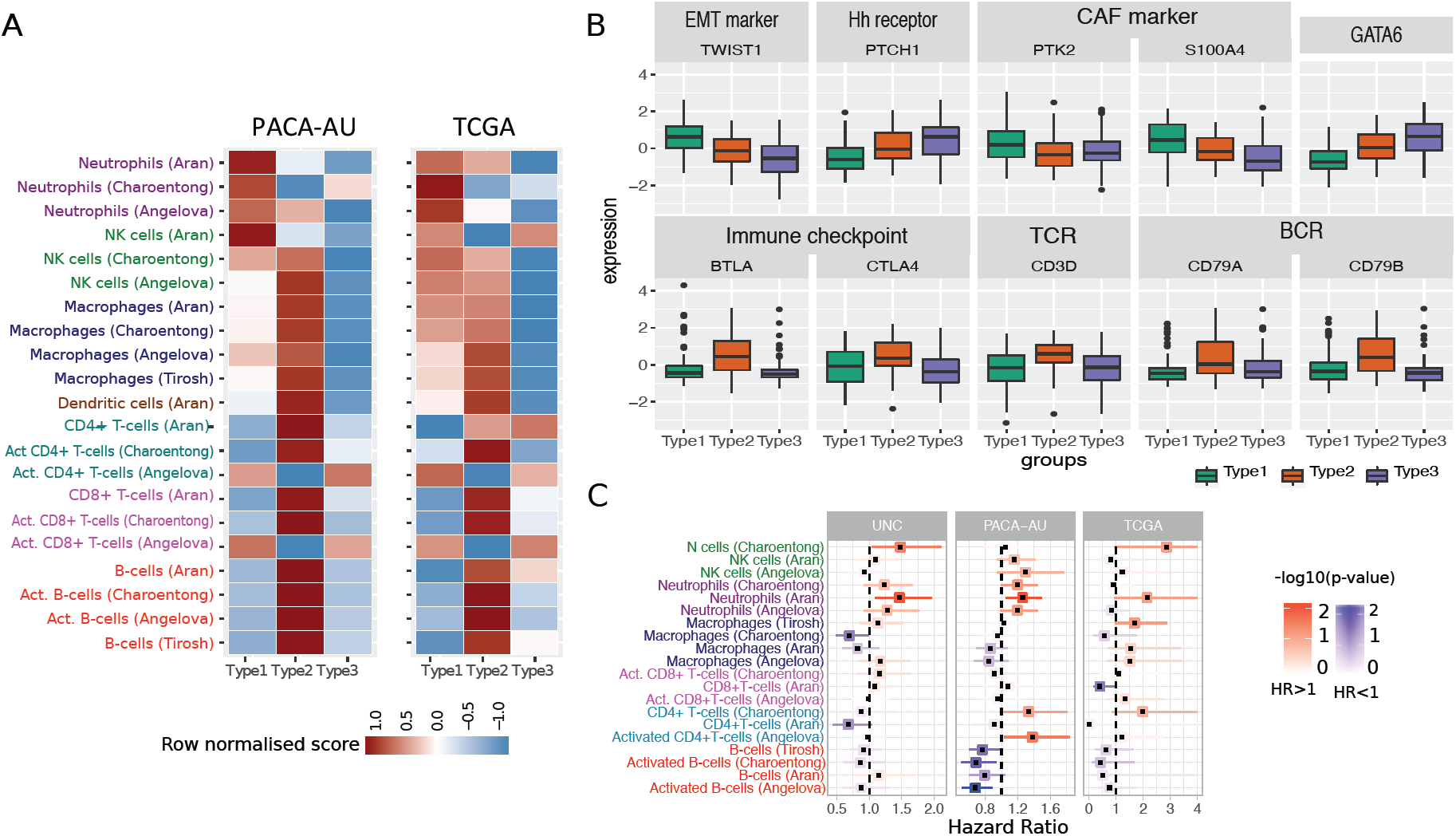
Tumour-infiltrating subpopulations of immune cells are associated with distinct PDAC subtypes. **(A)** Heat map of row scaled immune infiltrated scores per immune cell type. Angelova, Charoentong and Tirosh represent single-sample GSEA scores of signatures for cell types from the corresponding manuscripts. Aran is the inferences produced using xCell algorithm[21]. **(B)** Boxplots showing markers of special interest in PDAC subtypes differentially expressed between groups from the PACA-AU cohort. Similar trends were observed in the TCGA cohort (**Supplementary Figure S6**). EMT – Epithelial-mesenchymal transition; Eh – Hedgehog; CAF – Cancer Associated Fibroblasts; TCR – T-cell receptor; BCR – B-cell receptor. **(C)** Correlation of tumour infiltrating immune cells with patient overall survival. For three independent cohort (PACA-AU, TCGA and UNC), a multivariate Cox proportional hazards regression analysis was performed, with covariates including the enrichment scores of six immune cell types, and when available, patient gender, age at diagnosis, tumour stage and tumour grade. Squares represent the hazard ratio (HR) and the horizontal bars extend from the lower limit to the upper limit of the 95% confidence interval of the estimated of the hazard ratio. The colour scale reflects –log10(p-value) and is shown in blue for HR < 1 (good prognosis) and in red for HR > 1 (bad prognosis).

### Subtype 1: ‘innate immune’

Our subtype 1 consensus PDAC group showed an enrichment of Natural Killer (NK) cells and neutrophils, and an exclusion of other tumour infiltrated lymphocytes such as activated CD4+ T cells, CD8+ T cells and activated B-cells. Activation of primary drivers of EMT such as transforming growth factor-β (TGFβ) [6] and the Twist1 gene and de-regulation of developmental signalling pathways such as Hedgehog (Hh) and Wnt–β-catenin signalling [5, 7] have also been associated to tumours in this category by previous studies. We confirm such findings by showing marked up-regulation of Twist1 (EMT markers), and down-regulation of Ptch1 in samples of this subgroup (**Figure 3B;** Welch’s t-test FDR adjusted p-value < 10^-6^). Ptch1 is a receptor for Hedgehog (Hh) ligands and a tumour suppressor in the Hh pathway. Differential gene expression analysis also showed several de-regulated pathways related to ‘extracellular matrix organization’, ‘cell adhesion’ and ‘developmental processes’ (**Supplementary Table 4)**. The Wnt–β-catenin developmental pathway signalling, which was found to be up-regulated this PDAC group [7], is known to correlate with T-cell exclusion across solid tumours [25, 26]. This relationship was then validated recently in a clinical setting making the therapeutic strategy of beta-catenin inhibitors with immunotherapy a potential strategy for T-cell deficient tumours [27]. Bailey et al 2016 also identified gene programmes that included inflammation, hypoxia response and autophagy, and associated to the ‘squamous’ tumours, which are enriched in this group.

Type 1 tumours were associated with the worse survival (**Figure 2D,F**). Therefore, the desmoplastic stromal compartment and its interactions with tumour cells have clearly important roles in the poor outcomes for this group of PDAC tumours.

We performed multivariate survival analysis adjusting for several clinicopathological features, including tumour grade and tumour stage, and identified a negative relationship between neutrophil abundance and survival in PDAC (**Figure 3C**). We validate this prediction using three cohorts: PACA-AU, TCGA and an additional cohort of 102 PDAC tumour samples obtained from the University of North Carolina (UNC). The results of the enrichment of the immune infiltrates showed associations of neutrophils with survival in independent cohorts (**Figure 3C**) highlighting their potential as clinical biomarkers and therapeutic targets. This was also confirmed by an orthogonal method, the *TIMER* algorithm [28], a computational deconvolution method for inferring immune infiltrates, with neutrophils infiltration level significantly predicting patient survival (*P*=0.012 after adjusting for age, gender, ethnicity, stage and tumour purity; **Figure S5**).

### Subtype 2: ‘immunogenic’

The second PDAC cluster is characterized by PDAC tumours that displayed enrichment of many tumour infiltrating immune subpopulations related to adaptive immunity including activated CD8+ and CD4+ T-cells, and B-cells (**Figure 3A**). Subtype 2 samples exhibited a gene expression profile compatible with increased expression of genes associated with an ‘immune response’, ‘positive regulation of immune system process’ and ‘cell activation’ (**Supplementary Table 4**). This immune subtype of PDAC is characterized by marked upregulation of genes known to play roles in immune checkpoint inhibitions (e.g. CTLA4 and BTLA) [29]; B-cell receptor and T-cell receptor genes (e.g. CD3D, CD79A and CD79B) (**Figure 3B;** Welch’s t-test FDR adjusted p-value < 10^-2^). “Normal stroma”, ‘Exocrine-like’, ‘ADEX’ and ‘Notch’ PDAC tumour samples are also over-represented in this cluster (**Figure 1B**). Sivakumar et al., 2017 also observed enrichment for T cell–related pathways, such as those pertaining to T cell activation, proliferation, and differentiation, adaptive immune response and a significant prevalence of infiltrating CD8+ T cells in tumour samples enriched in this group.

Subtype 2 is the most immunogenic subtype with significant better survival when compared to samples of subtype 1. It has been shown before that higher levels of CD8+ T cell infiltration correlate with a better survival with Tumeh et al showing that CD8 T cell infiltration is needed for PD-1 therapy to work [8, 30, 31]. Together these findings indicate that this subtype is potentially amenable to therapy based on immune-check point inhibitors.

### Subtype 3: ‘Immune Exclusion’

Subtype 3 tumours exhibit lower enrichment scores for immune signatures, which suggests a lack of tumour-infiltrating lymphocytes in the microenvironment (**Figure 3A**). This group is enriched for PDAC tumours that have been characterised by high expression of adhesion-associated and epithelial genes [4] and genes with distinct roles in the control of cell-cycle, essential mitotic checkpoint functions, chromosomal stability, and DNA repair [7]. We confirmed that GATA6 gene expression is high in the subtype 3 group and low expression in subtype 1 (**Figure 3B;** Welch’s t-test FDR adjusted P < 10^-6^), which is expected given the overlap between subtype 3 and classical subgroups defined by Collison et al and Moffitt et al. Similarly, it has been recently shown that GATA6 expression inhibits the epithelial– mesenchymal transition (EMT) in vitro and cell dissemination in vivo and is associated to suppression of basal-like (like the one activated in subtype 1) molecular phenotype in PDAC tumours [32]. Collison et al. 2011, compared PDA cell lines representative of the classical and QM-PDA subtypes and described that the classical PDA cell lines are enriched in a KRAS-addiction gene expression signature and more dependent on KRAS than QM-PDA lines [4]. We found a strong enrichment for multiple metabolism signatures (**Supplementary Table 3**) in this group indicating prominent metabolic adaptation.

Moffitt et al., has shown that patients with ‘basal-like’ (subtype 1) tumours showed a strong trend toward better response to adjuvant therapy when compared to patients from the ‘classical’ (subtype 3) subtype group [5] and that QM-PDA cell lines were, on average, more sensitive to gemcitabine and less sensitive to erlotinib than the classical cell lines [5]. Additionally, it has been shown that patients with GATA6^high^ PDAC tumours have better response to 5-FU when compared to ‘basal-like’ GATA6^low^ patients [32]. Together these results suggest that *KRAS*-directed therapies or therapies targeting growth pathways such as EGFR-targeted therapy might be best deployed in this subtype 3 classical PDAC subtype when compared to the other PDAC subtypes.

### Mutational signature analysis

Multiple mutation signatures have been established through pan-cancer analysis of cancer genomes [33]. These have established mutational signatures for pancreatic cancer including a signature associated with failure of double strand break repair by homologous recombination (Signature 3) [33]. Using a panel of thirty mutational signatures from COSMIC, we established the contribution of each signature to the cohort of pancreatic cancers and sought to identify if any signature that is enriched for the immunophenotypes we have described in this manuscript. We found three major contributory signatures in these pancreatic cancers (**Figure 4A** and **Figure S7**) but none of these were enriched for a particular subtype. These were signatures 1, 6 and 15 [33]. Signature 1 correlates with the aging process and 6 and 15 are tied to DNA repair. The signature analysis does not help explain the difference in subtypes but does reinforce that certain cases of pancreatic cancer can benefit from treatment with therapeutics against DNA repair.

**Figure 4.**
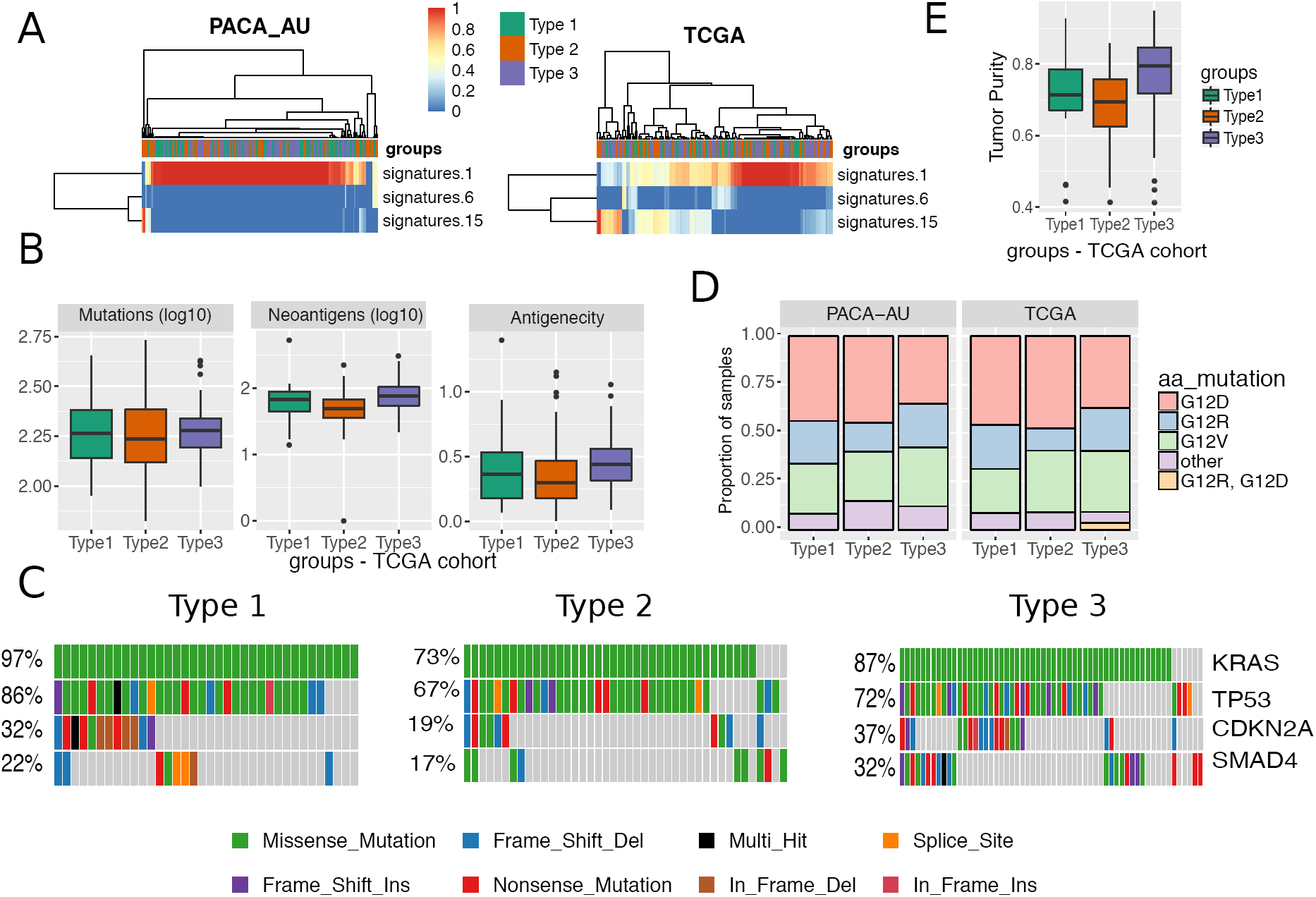
Mutation analysis. **(A)** Heatmap showing the relative contribution of mutation signatures 1, 6 and 15. **(B)** Mutation load, neoantigen load and antigenicity for different tumour subtypes of the TCGA cohort **(C)** OncoPrint displaying frequency of mutated genes in different PDAC subtypes of the TCGA cohort **(D)** Distribution of activating KRAS mutations across the three PDAC subtypes in the TCGA and PACA-AU cohorts (E) Tumour purity (ESTIMATE scores) across subtypes of the TCGA cohort.

### Mutational burden

Evidence suggests that the degree of mutagenesis within a tumour may represent a biomarker for response to immunotherapy. It is thought that highly mutated tumours are more likely to harbour neo-antigens, which make them targets for adaptive immunity. Tumour mutation burden has been shown, in several tumour types, to correlate with patient response to both CTLA-4 and PD-1 inhibition [34, 35]. Similarly to Charoentong et al., 2017, we defined the frequency of neoantigens (i.e. number of neoantigens / number of mutations) in a given sample as a surrogate of tumour antigenicity. We found a significant difference between antigenicity in subtype 2, which was lower when compared to subtype 3 (Wilcoxon test P=0.003; **Figure 4B**). The most frequently mutated pancreatic cancer genes (KRAS, CDKN2A, SMAD4 and TP53) are equally distributed across the three subtypes (Figure 4C), except for KRAS and SMAD4, which are observed in lower frequency in subtype 2 (73% and 17%, respectively) when compared to subtype 1 (KRAS: 97%; Fisher exact test *P* = 0.003) and subtype 3 (SMAD4: 32%; Fisher exact test *P* = 0.03). We also analysed the status of the activating KRAS mutation, namely G12D, G12R, G12V and other (A11T, G12A, G12C, G12L, G12S, G13C, G13P, GQ60GK, Q61H and Q61R) and verified that KRAS mutations are spread out equally over the three subtypes in both cohorts (Fisher exact test P >> 0.05; **Figure 4D**). In summary, smaller frequency of neoantigens of key pancreatic cancer driver mutations are observed in the subtype 2 PDAC subtypes. However these results are inconclusive because it is highly possible that the level of purity across tumour samples affects the interpretation of genomic analyses [36]. In fact, we observed striking differences in estimated promotion of tumour cells in the samples between PDAC subtypes, in particular subtype 2 displayed the lowest purity (**Figure 4E; Supplementary Figure S9**). This variability confounds the interpretation of genomic analysis results when comparing between PDAC subtypes.

## Discussion

In this study, we propose a consensus clustering of PDAC into three major subtypes that have different molecular and clinical characteristics and may respond differently to selected therapies. We suggest that PDAC subtypes should be considered when stroma and immune modulating therapies are studied in the future. The identified consensus applies to tumour samples from four independent cohorts, demonstrating the robust nature of this new subtype classification in PDAC. It is important to notice the strong presence of stromal components in subtypes 1 and 2, but also distinct prognosis for each group. However, while the subtype 1 group of tumours is characterised by a reactive desmoplastic stroma and an inflammatory microenvironment with possible epithelial-to-mesenchymal transition (EMT) events; the stromal compartment in subtype 2 is enriched in infiltrated CD8+ and CD4+ T-cells.

Subtype 1 suggests that the functional role of EMT regulators and innate immune cells in immune evasion is complex. The biological link between the inflamed immune subtype and EMT is consistent with the finding that the stroma of subtype 1 tumours is infiltrated not only with innate immune cells, but also markers typically found in activated cancer-associated-fibroblasts (e.g. PTK2 and S100A4; Welch’s t-test FDR adjusted p-value = 0.02 and 3x10^-6^, respectively; **Figure 3B**). In addition, it suggests that the worse outcomes seen in the subtype 1 may be partially linked to a pro-metastatic immune evasive microenvironment. Tumour samples in this group have been characterized by infiltration of desmoplastic stroma (‘activated-stroma’, Moffitt et al., 2015) and high expression of mesenchyme associated genes (‘basal-like’, Collison et al., 2011). These results corroborate initial findings by Guerra et al., and others that inflammation increases increase both EMT and cancer cell invasion [21, 22] and that the presence of IL-6 pro-inflammatory marker in the serum of patients with pancreatic cancer has been associated with worse survival [23]. A better definition of the tumour-expressed ligands recognized by these myeloid cell subsets and their role in driving tumour progression and antitumour immunity will facilitate more detailed functional analyses and identify possibilities for therapeutic intervention.

Another challenge that arises is the definition of better preclinical models that recapitulate these subtypes. The interplay between the stroma and the immune components are difficult to model. Previous research suggests that traditional pancreatic cancer cell lines from the Broad Institute Cancer Cell Line Encyclopaedia only recapitulate the ‘classical’ subtype [7] and PDX models are not able to recapitulate the ‘normal-like’ subtype [5].

Current tumour models such as the KPC mice do not accurately reflect variations in sub-types of pancreatic cancer and there is generally difficulty at obtaining high quality primary samples. Less then 20% of patients undergo a resection and most of the tumour is filtrated with a desmoplastic stromal reaction that composes of collagen, fibroblasts and immune cells. Studies so far have tried to enrich as best as possible the tumour compartment including the ones we have used. In our study we have been able to identify the distinct characteristics of immune components from RNA sequencing of pancreatic tumours and we have described the possible clinical significance of these findings. Our findings further suggest that characterisation of the non-tumour components of pancreatic cancer will be revealing. Single cell sequencing of primary tumours will enable improved understanding of the complex interplay in the stromal-tumour microenvironment.

Several lines of evidence suggest differential drug response sensitivity between the different subtypes. The regulatory contribution of the immune system should be assessed more thoroughly in human PDAC cancer to guide new therapeutic interventions tailored to patients with different tumour subtypes. Understanding PDAC subtypes could be used improve better patient stratification for clinical drug trial enrichment schemes to better select patients to make detection of a treatment effect more likely. However, more detailed immune characterization of PDAC tumours is needed.

## Methods

### Data download: Gene expression profiles, clinical and mutations datasets

The PACA-AU gene expression data (*n* = 269) plus clinical and mutational profiles were obtained from ICGC data portal (https://dcc.icgc.org/releases/release_24/Projects/PACA-AU). TCGA ‘*htseq.count.gz*’ raw counts files for RNA-seq data (*n* = 177) were obtained from the GDC portal (https://portal.gdc.cancer.gov/). Clinical metadata and Mutation annotation files (MAF) for the TCGA cohort was obtained from the GDC legacy archive (https://portal.gdc.cancer.gov/legacy-archive/) with the UUIDs: a9f29dc4-6a6a-42f3-b06d-9e6ded926b55 (clinical metadata) and faf50bd9-bfc8-4dfa-b0ca-9184e44fb07f (MAF file). The UNC gene expression data (n = 132) plus clinical profiles were obtained from gene expression omnibus (GEO) archive under the accession number GSE21501. Out of 132, in the UNC cohort, 30 were excluded due to unavailability of survival time in the clinical table.

### Processing of gene expression data

Batch effects were removed by applying the ComBat algorithm [37]. TCGA RNA-seq data was transformed with the variance-stabilizing transformation method [38] prior to removing batch effects. Batch IDs for the TCGA cohorts were obtained from the sample barcode, namely the 'plate' id as described in https://wiki.nci.nih.gov/display/TCGA/TCGA+Barcode.

### Classification of PACA-AU and TCGA cohorts according to five previous classification schemes

The R package ConsensusClusterPlus [39] was employed to subtype PDAC samples according to the expression signatures defined in Moffitt *et al.* [5] and Collisson *et al* [4] (**Supplementary Figure S1**). The number of clusters was confirmed by examining cumulative distribution function (CDF). We confirmed the existence of well separated clusters for Moffitt *et al.* classification based on tumour (two clusters: basal-like and classical) and stroma signatures (2 clusters: stroma and activated stroma). For the Collison *et al.* classification we confirmed the existence of evident 3 clusters (classical, exocrine-line and quasimesenchymal). Bailey et al cluster labels were directly downloaded from (PACA-AU cohort) [6] and TCGA cohort) [40]. Sivakumar et al cluster labels were directly downloaded from [7].

### Identification of consensus subtypes

To identify a meaningful consensus of the multiple clustering labels we used consensus clustering and the partition around medoids (PAM) algorithm to cluster the PACA-AU and TCGA cohorts according the similarity of their labels. For the PACA-AU cohort, we only included tumours that were classified as “Pancreatic Ductal Adenocarcinoma” according to the tumour histological type classifications and were also labeled by all 5 classification schemes which corresponded to 204 PDAC tumours. For the TCGA cohort we included only the filtered PDAC cases according to *Raphael et al*, this corresponded to 149 TCGA tumours [40]. The Hamming distance was used as a measure of similarity between PDAC tumours. The robustness of sample classification was analysed by examining cumulative distribution function (CDF) of the proportion of times in which 2 samples are clustered together across the resampling iterations (×1000) [41]. For varying number of clusters (K=2 to K=7), we examined the area under the curve of the consensus distribution function (CDF) plot (**Supplementary Figure S2 and S3**) and identified three robust consensus subtypes in both cohorts.

### Survival analysis

Multivariate Cox regression, log-rank test and Kaplan–Meier estimators were implemented using the R package survival. For the PACA-AU cohort (**Figure 2C**), we adjusted the survival differences for age, gender, and tumour stage, tumour grade and tumour relapse covariates. For the TCGA cohort (**Figure 2E**), we adjusted for age, gender, tumour stage, tumour grade, new event indicator (yes/no) and targeted therapy indicator (yes/no). Tumour stage refers to the TNM Staging System based on the extent of the tumour (T). The targeted therapy indicator refers to whether the patient had adjuvant and/or postoperative pharmaceutical therapy. More specific information about the treatment regime was not possible to obtain. The correlation between immune cell scores and survival (**Figure 3C**) was performed using a multivariate cox regression model adjusting for age, gender, tumour grade and stage and tumour relapse (PACA-AU and TCGA cohorts) and adjusting for tumour stage (TMN T stage) for the UNC cohort. For the TCGA cohort we only included cases with no targeted therapy.

### Analysis of immune infiltrates and tumour purity

Two main methodologies were used to identify immune cell types enriched in the tumour microenvironment. For three of the gene signatures [22-24] we used single-sample gene set enrichment analysis (ssGSEA) method implemented in the GSVA R package[42]. For the Aran gene signatures we used the xCell method implemented in R [21]. The xCell method also relies on ssGSEA analysis but contains an additional step which uses a reference matrix of ‘spillovers’ between cell types. The ‘spillover’ step is thought to better eliminate dependencies between closely related cell types (e.g. such as between CD8+ T-cells and NK cells). Log2(TPM+1) gene expression levels from the TCGA cohort were used to estimate the enrichment with the ssGSEA method, and TPM levels were used with the xCell method. Normalised microarray gene expression levels (Illumina Expression BeadChIPs) from the PACA-AU cohort were used directly in both methods.

ESTIMATE was used to gauge the degree of leukocyte infiltration, stromal content and tumour purity[2]. Data was summarised per PDAC subtypes in the PACA-AU and TCGA cohorts (**Figure 4E; Supplementary Figure S6**)

### Mutation analysis

Mutational signature analysis was performed using the DeconstructSigs R Package [43]. We determined the contribution of thirty signatures defined in COSMIC (http://cancer.sanger.ac.uk/signatures/) to explain each pancreatic mutational profile. Normalisation was relative to the number of times each trinucleotide context is observed in the exome. The output was a set of weights specifying the estimated contribution of each of the 30 known signatures to the mutation profile. MAF files were parsed with maftools [44] (**Supplementary Figure S8**). Number of neoantigens per tumour in the TCGA cohort was retrieved from The Cancer Immunome Atlas database (https://tcia.at/home) [22].

### Statistical analysis

All statistical analyses, Fisher’s exact test, Chi-square test, Wilcoxon rank sum test, hypergeometric test, and hierarchical clustering, were performed using R [45]. Multiple test correction was performed using the R function p.adjust and the Benjamini and Hochberg (FDR) method. Jaccard coefficients were computed using the R package arules. Differential expression analysis between the groups was carried out using the Welch’s test implementation in R and by comparing each group against all others. Welch’s test is a variant of the classical Student test, whose goal is to test the equality between two means taking assuming different variances between two groups. When necessary, Ensembl or Entrez IDs were mapped to human HUGO identifiers using Ensembl version 89 biomart (http://www.ensembl.org/biomart)

